# Electromagnetic Field and Temozolomide Increase Differentiation of Human Glioblastoma Cell Line; concise view of mechanism

**DOI:** 10.1101/340596

**Authors:** Meysam Ahmadi-Zeidabadi, Zeinab Akbarnejad, Marzie Esmaeeli, Yaser Masoumi-Ardakani, Hossein Eskandary

**Author notes:** These authors contributed equally to this work (Meysam Ahmadi-Zeidabadi, Zeinab Akbarnejad). Corresponding author at: Neuroscience Research Center, Kerman University of Medical Sciences, 76175-113, Kerman, Iran. Email addresses (H. Eskandary).

## Abstract

Glioblastoma is a highly malignant brain tumor with an extremely dismal prognosis, with a median survival of 12 months. Promoting glioma stem cell (GSC) differentiation is a crucial therapeutic strategy in treating glioblastoma to improve survival. We combine standard chemotherapy drug Temozolomide (TMZ) with Electromagnetic Field to evaluate their differentiation effects on glioma U87 cell line.

Human glioma U87 cells exposed to electromagnetic field (EMF), Temozolomide (TMZ) alone and a combination of both were compared to control group. Nestin and CD133 were detected to identify stem-like cells (SLCs) changes during the experiment. The differentiation was found through detecting the expression of the glial fibrillary acidic protein (GFAP), and Notch4. We evaluated Ca^+^, SOD, and Notch as important chemical mediators in signal transduction to elucidate the mechanism of actions in the processes of differentiation.

The expression of cancer SLC markers CD133, and neural stem/progenitor Nestin decreased in EMF/ Drug combined group compared to control which shows depletion of stem-like cell pool. In contrast, the differentiation of SLCs was increased by detecting the expression of the glial fibrillary acidic protein (GFAP), but, Notch4 decreased in Electromagnetic field/Drug combined group compared to control which shows a benign process of treatment. The differentiation was also confirmed by light microscopy. Morphological changes such as an elongated shape with elongated neurites, intercellular connection, and neurite branching were observed. Superoxide dismutase (SOD), Ca^+^ and Notch were also increased as important chemical mediators in signal transduction.

Here, we found the combination of pulsed electromagnetic fields (PEMFs) and TMZ significantly resulted in differentiation and glioma stem cells (GSCs) pool reduction that could have a profound therapeutic implication.

## Introduction

Glioblastoma multiform (GBM) is the most common and aggressive form of primary brain tumor in adults. Despite standard treatment with surgery, irradiation, and chemotherapy to extend survival, it universally recurs and unrelentingly results in death. Their ability to infiltrate diffusely into the normal brain parenchyma is associated with the worst prognoses [1]. The median survival is approximately 12 months with less than 10% of patients alive at 5 years after diagnosis [2]. This dismal prognosis of GBM has motivated researchers to investigate alternative therapeutic paradigms, such as differentiation therapy. The vast majority of research has been focused on the aspects of proliferation and apoptosis, and little is known about its possible role involved in the process of cancer cell differentiation.

It has recently been accepted that undifferentiated tumor cells, called cancer stem cells (CSCs) play a pivotal role in the initiation and progression of cancers in various tissues [3]. CSCs comprise only a small portion of the tumor, and every single cell can give rise to a new tumor. If the malignant cells of cancers are cancer stem cells, then it should be possible to treat cancers by inducing differentiation of the stem cells. If tumor cells can be forced to differentiate and cease proliferation, then their malignant potential will be controlled [4–6]. Increasing evidence indicates that malignant gliomas originate from neural stem-cell populations, which are termed glioma stem-like glioma cell [7,8]

Currently, drug-induced differentiation is considered as a promising approach to eradicating tumor-initiating cell population and some of this anticancer drugs appeared to have the capability of inducing glioma cell differentiation [9,10].

The addition of Temozolomide to radiotherapy for treatment of newly diagnosed glioblastoma patients increases median survival 2 -2.5 months [11,12]. The study of Yuan et al., showed that Resveratrol and TMZ significantly resulted in an increase in expression of astrocyte differentiation marker, glial fibrillary acid protein (GFAP). In their study, TMZ treated group showed a significant GFAP expression compared to control group [13]. Beier et al., reported, Temozolomide induced a dose and time-dependent decline of stem cell subpopulation from MGMT negative tumor [14].

Pulse electromagnetic field in the 0-300 H_z_ range as a therapeutic tool extensively used for the treatment of cancer [15,16]. Some studies have suggested that static magnetic fields (SMFs) and pulse electromagnetic fields have the potential as the adjunctive treatment method for chemotherapy [17–19]. Previous studies have demonstrated that the cellular effects of extremely pulsed electromagnetic fields (ELF-EMFs) depend, in large part, on their intensity and exposure time, as well as on the phenotype of the cellular target and interactions with intracellular structure [1 7,20,21]. Application of different time course of ELF-EMF have been shown to induces a different pattern of differentiation irrespective of co-stressor applied [20]. Also, electromagnetic field has been reported to induce cell differentiation of stem cells [22]. Some studies illustrate the potential of EMF therapy in combination with conventional cancer therapies to sensitize tumors [23].

Improvement of glioblastoma treatment by even a moderate amount could potentially result in many years of life saved. Differentiation-inducing therapy, which modifies cancer cell differentiation, has been proposed to be a novel potential approach to treat malignant tumors. As mentioned above TMZ and EMF have the abilities of differentiation and decreasing of SLGC or GBM-CSCs that express certain stem cell-associated markers, including CD133 [24].

Although recent observations have shown that tumorigenicity is not entirely restricted to CD 133 positive tumor subpopulation [24], there is a large, growing set of data linking the expression of AC133/CD133 with poor patient prognosis in glioblastoma multiform patients [25]. AC133/CD133 downregulation is reported to be associated with better survival in a mouse model [5]. Some data indicate that tumorigenicity, as well as radio-resistance and chemo-resistance, may be attributes of CD133 expressing SLGCs [4,26]. If eradication of SLGCs is the critical determinant in achieving cure [27], it must be reasoned that depletion of the AC133/CD133-positive cell pool through controlled, agent or drug-induced differentiation could have profound therapeutic implications. Inducing differentiation is one of the crucial therapeutic methods in cancer treatment [13]. The purpose of this study was to evaluate the effects of TMZ and PEMFs on differentiation and GSCs reduction.

To confirm this assumption, we studied the effect of TMZ and PEMFs on glioma cell line U87. We exposed human glioma U87 cells to three groups of experiment including; EMF, TMZ alone, and combined treatment, and compared them to control group. The expression of cancer SLC markers CD133 and neural stem/progenitor Nestin were detected to identify SLCs. And GFAP and Notch was assessed for differentiation of SLCs. We evaluated the Ca^+^, SOD, Notch and some important chemical mediators in signal transduction to elucidate the mechanism of actions in the processes of differentiation. Any therapeutic agent that induces differentiation and deplete stem-like cell pool, could have profound therapeutic implications. Here, we found that the combination of PEMFS and TMZ significantly resulted in differentiation and GSCs reduction pool.

## Material and Method

### Cell culture

Human glioblastoma cells from the U87 cell line (Pasteur Institute, Tehran, Iran) were cultured in 25 cm^2^ flasks (Iwaki, Tokyo, Japan) at a cell density of 10^4^ cells/ml in Dulbecco’s modified Eagle’s medium (DMEM; product code 11966025-Invitrogen Inc. Gibco BRL, Gaithersburg, MD, USA). Supplemented with 10% (v/v) heat inactivated, sterile-filtered, fetal calf serum (Product Code: 12306C-500ML-Sigma-Aldrich Co., St. Louis, MO, USA) and 1% (v/v) of penicillin-streptomycin solution (Product Code: P4333-100ML-Sigma-aldrich Co.), in a humidified atmosphere of 5% CO2 at 37 ± 0.5 °C. The cells were continuously exposed to different EMF (100 Hz, 100 ± 20 G), TMZ (100 μ M) and (EMF and TMZ) up to 144h. Control cells remained unexposed to EMF and TMZ for the same time.

### Exposure system

The exposure system has been described in greater detail elsewhere [17]. Briefly, ELF-EMF was generated by a clinically approved magnetotherapy device Fisiofield Mini (Fisioline Co., Verduno, Italy) that could generate square or sinusoidal waves with frequencies of 0–100 Hz and amplitudes of 0 – 100 G. The device was set to generate continuous square waves with a frequency of 100 Hz and amplitude of 100 G. In addition, the device contained four coils that could apply magnetic fields to two groups as simultaneously and individually. As shown in Fig. 1A and B, each culture flask was placed between two identical coils, each coil put in a chamber of Plexiglas sized 160 Å~ 160 Å~ 50 mm, by maintaining the coaxial distance of 40 mm apart. A cooling system based on continuous recirculation of cold water around each coil prevented the temperature from rising. All materials used to build the exposure system (i.e., plexiglas, polystyrene, and PVC) were transparent to EMF. The magnetic amplitude measurements, by including the amplitude values generated by the device, were made by means of a Gaussmeter GM08 (Hirst Magnetic Instruments Ltd., Falmouth, UK) equipped with a Hall-effect sensor. (Fig. 1)

**Fig.1.**
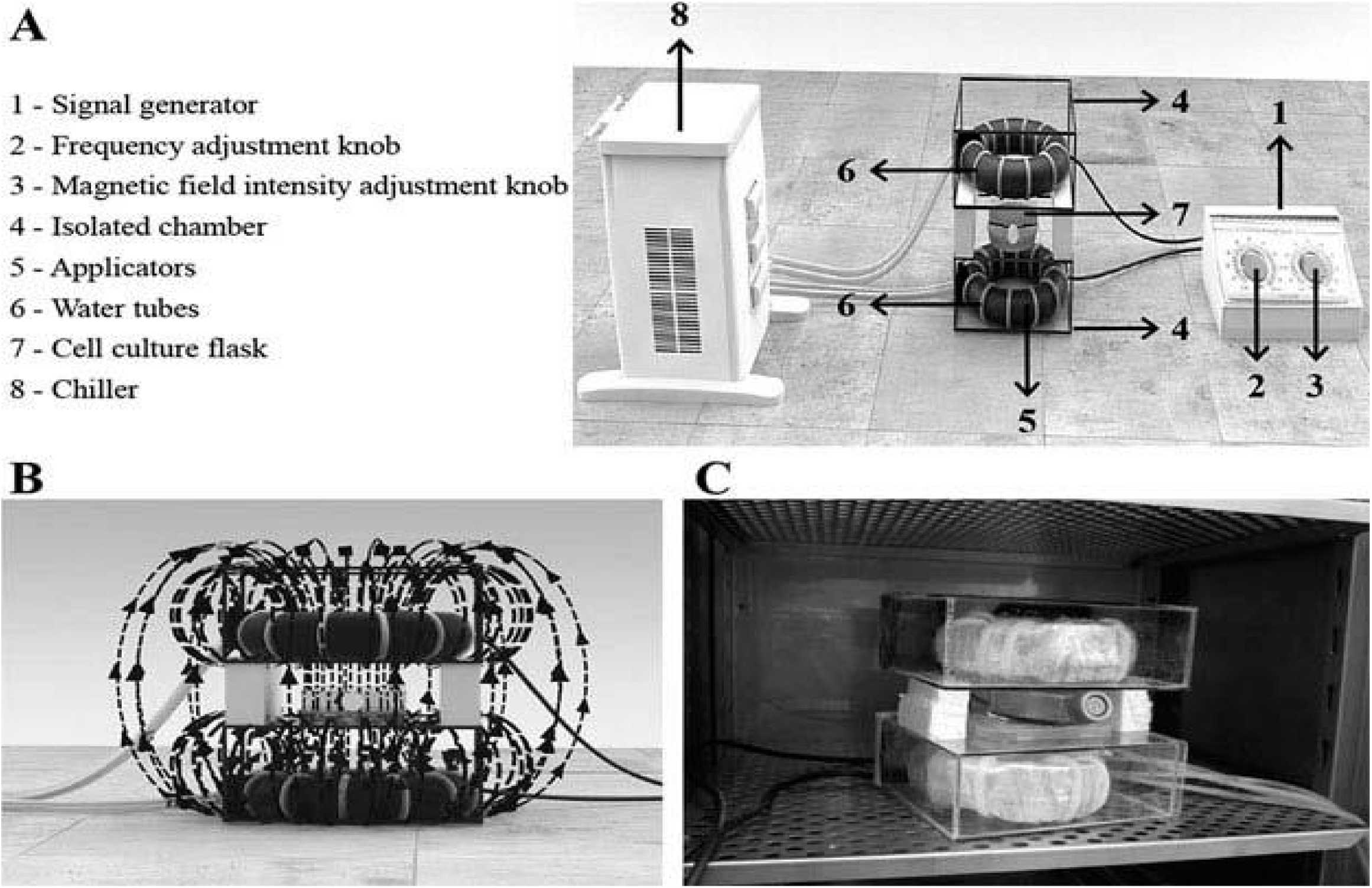
ELF-PEMFs exposure system. A) The cell cultures were exposed by using a house-made ELFPEMFs exposure system constituted of: (1) a signal generator Fisiofield Mini (Fisioline Co, Verduno, Italy) set to generate continuous square waves with a frequency of 10, 50 or 100 Hz and amplitude of 50 or 100 G, which are selectable by using apposite knobs (2-3); (4) two chambers made of plexiglas sized 160×160×50 mm, each one containing a copper coil wire with a diameter of 130 mm (5). The two chambers were separated by two side thicknesses of polystyrene, maintaining a coaxial distance of 40 mm, in order to allow the housing of a culture flask (6); a water-based cooling system constituted of delivery tubes (7) and a chiller (8). B) The part of ELFPEMFs exposure system including coils and culture flask has been designed in order to fit inside the incubator. C) not in scale magnetic flux showing the direction of the field lines.

### SOD assay or Measurement of intracellular reactive oxygen species (ROS) formation

Superoxide dismutase (SOD) is an antioxidant enzyme involved in the defense system against reactive oxygen species (ROS). SOD level was measured using enzyme-linked immunosorbent assay (ELISA) kit (ABNOVA, Taiwan). Briefly, after performance of the treatments, the cells were separated by EDTA trypsin and centrifuged for 20 minutes (at 3000 rpm) then supernatants were carefully collected. According to the manufacture’s recommendations (bioassay technology laboratory) dilution of standard solutions performed. For standard well, 50 μl of standard, 50 μl Streptavidin-HRP solution (since the standard has already combined biotin antibody, it is not necessary to add the antibody) and for test well, 40 μl of sample, 10 μ l of SOD antibody and 50 μl of Streptavidin-HRP were added. Then seal the sealing membrane, gently shacked and incubated 60 minutes at 37° after washing with drain liquid, chromogenic solution A 50 μl, chromogenic solution B 50 μl to each well was added. After mixing, incubate for 10 min at 37° away from light for color development. Finally, 50 μ l of Stop solution was added into each well to stop the reaction (the blue changes into yellow immediately). Then color progresses between 10-20 min were assessed by ELISA reader (450nm, reference wavelength 630nm).

## Determination of intracellular calcium concentration

Measurement of intracellular Ca^2+^ Calcium concentration is useful to study the upstream and downstream events of Ca^2^+ signaling. Different groups of cells were loaded with 5μM Fura 2-AM for 60 min. After washing and removal of Fura 2-AM, the cultures were incubated in dye-free saline for at least 45 min to allow for the de-esterification of the Fura2-AM. The cells were analyzed immediately on the fluorescence plate reader (FLX8oo, BioTek. USA) and the fluorescence intensity of cells in 96-well plates was quantified at an excitation of 340/380 nm and an emission of 510 nm. Each experiment was performed six independent times. Results were expressed as fluorescence percentage of control cells.

### MDA measurement

Malondialdehyde (MDA) is an index of peroxidation. The MDA in the sample is reached with Thiobarbituric Acid(TBA) to generate the MDA-TBA adduct. Briefly, 1×10^6^ cells were homogenized with 300μl of the MDA Lysis Buffer, then centrifuged (13,000 xg,10 min) to remove insoluble material. Place 200μl of the supernatant from each homogenized sample into a micro centrifuge tube. According to the manufacturer’s recommendations (Bioassay Technology Laboratory) dilution of standard solutions performed. Then, added 600μl of TBA solution into each vial containing standard and sample. Incubate at 95°C for 60 min. Cool to room temperature in an ice bath for 10 min. Pipette 200μl (from each 800μl reaction mixture) into a 96-well microplate for analysis. The optic density (OD) was then read at 532nm by enzyme-linked immunosorbent assay (ELISA) reader (Pharmacia Biotech, Stockholm, Sweden).

### Real Time PCR

Gene expression of (CD133, N) were analyzed by real-time polymerase chain reaction (PCR). RNA was extracted from 10^4^ cells by the guanidine isothiocyanate-phenol-chloroform method using RNX+ reagent (Cinagen, Tehran, Iran) [28]. The isolated RNA was solved in 20 ml of RNase-free water. The single-strand cDNAs were synthesized from total purified RNA using M-MuLV reverse transcriptase and oligo (dT) primer (Fermentas, Vilnius, Lithuania). Real-time PCR reactions were done using Step One device (Applied Biosystems Foster City, CA, USA) by mixing 10 ml of qPCR Master-MixSyGreen (Analytik Jena AG, Jena, Thuringia, Germany), 0.5 ml of primer forward, 0.5 ml of primer reverse, 2 ml of cDNA, and 7 ml of RNase-free water. PCR amplification (40 cycles) was performed in the following way: initial denaturation for 3 min at 95°C, denaturation for 30 s at 95°C, annealing for 30 s at different temperature according to primer sequences (table 1), extension for 40 s at 72°C and final extension for 12 min at 72°C. Primer sequences (table1). The gene expression quantification was done by using the 2^-ΔΔct^ method [29] and the REST 2009 software (QIAGEN, Hilden, Germany)

**Table.1.**
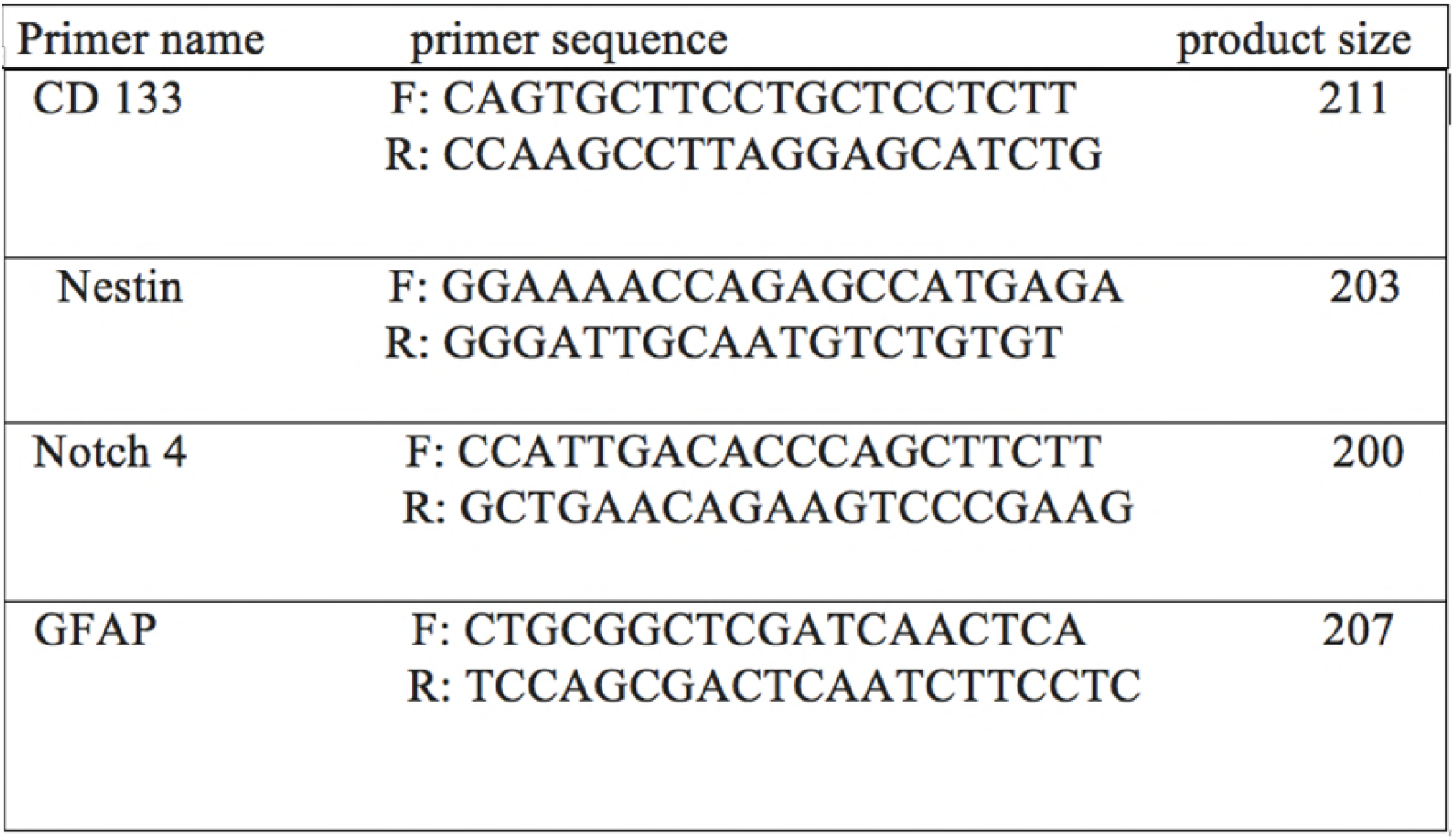
The nucleotide sequences that were used in primers for real-time analysis

## Western-blot analyzing

GFAP expression analysis by Western blots; Western blots of GFAP protein was carried out in order to study differentiation progression. 25 mg of total protein from cell lysate was separated by sodium dodecyl sulphate-polyacrylamide gel electrophoresis (SDS-PAGE) according to Laemmli [30] and then electro-transferred onto polyvinylidene fluoride (PVDF) membrane (Merck Millipore, Billerica, MA, USA), according to Towbin et al. [31]. After blockage for 2h at RT with 5% (v/v) instant non-fat dry milk/Tris-buffered saline solution containing 0.1% (v/v) Tween 20 (TBS-T), PVDF membrane was incubated overnight at 4°C with the primary antibody against GFAP 56055 – Santa Cruz Biotechnology Inc., Dallas, TX, USA). Specific antibody binding was detected by incubating the membranes with horse radish-peroxidase (HRP)-conjugated secondary antibody (Product Code: sc-2005 – Santa Cruz Biotechnology Inc.) at RT for 2 h. Luminol (GE Healthcare Bio-Sciences Corp., Piscataway Township, NJ, USA) was used as a chemiluminescent substrate for HRP. The membranes were washed with TBS-T and scanned by ChemiDocTM XRS+ imaging system (Bio-Rad, Hercules, CA, USA). OD-based quantification was performed by Image LabTM 3.0 (Bio-Rad). b-actin (Product Code: sc-81178 _ Santa Cruz Biotechnology Inc.) Western blots were used as controls.

### Light microscopy (LM)

An inverted phase-contrast microscope Axiovert 25 (Carl Zeiss AG, Oberkochen, Germany) was used to investigate morphological cellular changes. Three randomly selected fields from each culture were captured.

### Statistical analysis

The results are expressed as mea± SEM. The difference in mean intracellular ROS and calcium, MDA and genes and protein expression data, between experimental groups and control were determined by one-way ANOVA, followed by Tukey. P<0.05 was considered statistically significant. Data analysis was done by using statistical package for social science (SPSS) version 16 software (IBM, Armonk, NY, USA).

## RESULTS

### SOD activity

SOD activity increased after exposure of EMF and TMZ alone and concomitantly. SOD activity indirectly represents the ROS production. EMF, TMZ administration and combined treatment (EMF+ TMZ) increased activity of SOD 47%,60% and 85%after 120h and about 48%,75% and 98% after 144h respectively in comparing to control (Fig2).

**Fig. 2.**
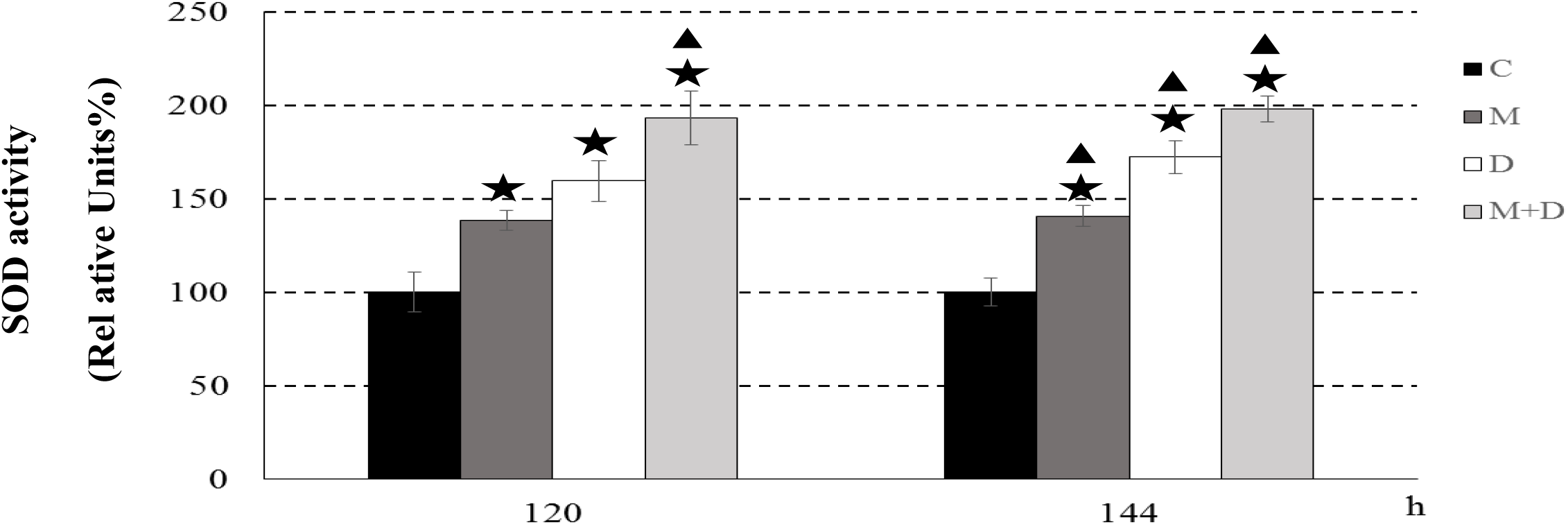
SOD activity was assessed. Values are reported as percentage of the control untreated cells, by considering it as 100%. Each value represents the mean ± SEM of four independent experiments, each done in duplicate. Star indicates the value significantly different from the control untreated cells (p < 0.05). Triangle indicates value significantly different from all the other treatments at the same time point (p < 0.05).

### Calcium concentration

Calcium concentration was analyzed by Fura-2 assay. Data showed, changes in calcium concentration level after 120h and 144h. Exposure of EMF and TMZ alone and concomitantly increased calcium level about 35%, no change and 75%after120h. Also, EMF, TMZ and co-administration increased calcium level about 40%,10% and 80% respectively after 144h (fig.3).

**Fig.3.**
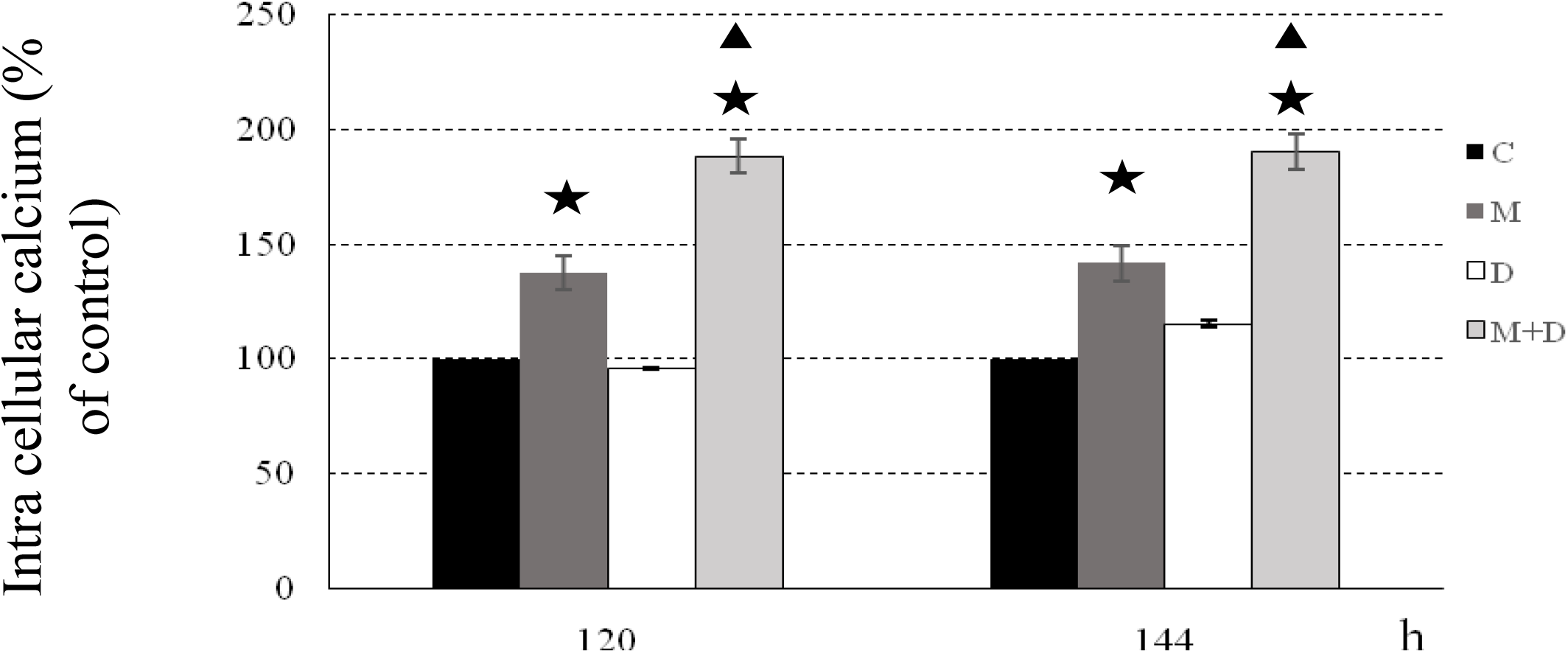
Determination of intracellular calcium of cells from different groups. Values are reported as percentage of the control untreated cells, by considering it as 100%. Each value represents the mean ± SEM of four independent experiments, each done in duplicate. Star indicates the value significantly different from the control untreated cells (p < 0.05). Triangle indicates value significantly different from all the other treatments at the same time point (p < 0.05).

### MDA measurement

According to our data lipid peroxidation increase significantly after treatment (EMF, TMZ) and co-treatment (EMF+TMZ) had synergistic effect. Malondialdehyde (MDA) value increased after 120h and 144h after exposure of EMF and TMZ alone and concomitantly. EMF increased value of MDA 2.3, and 6 folds after EMF, TMZ, and combination at 120h and 2.5,3.5 and 6.5 folds after 144h, comparing to control (fig.4).

**Fig.4.**
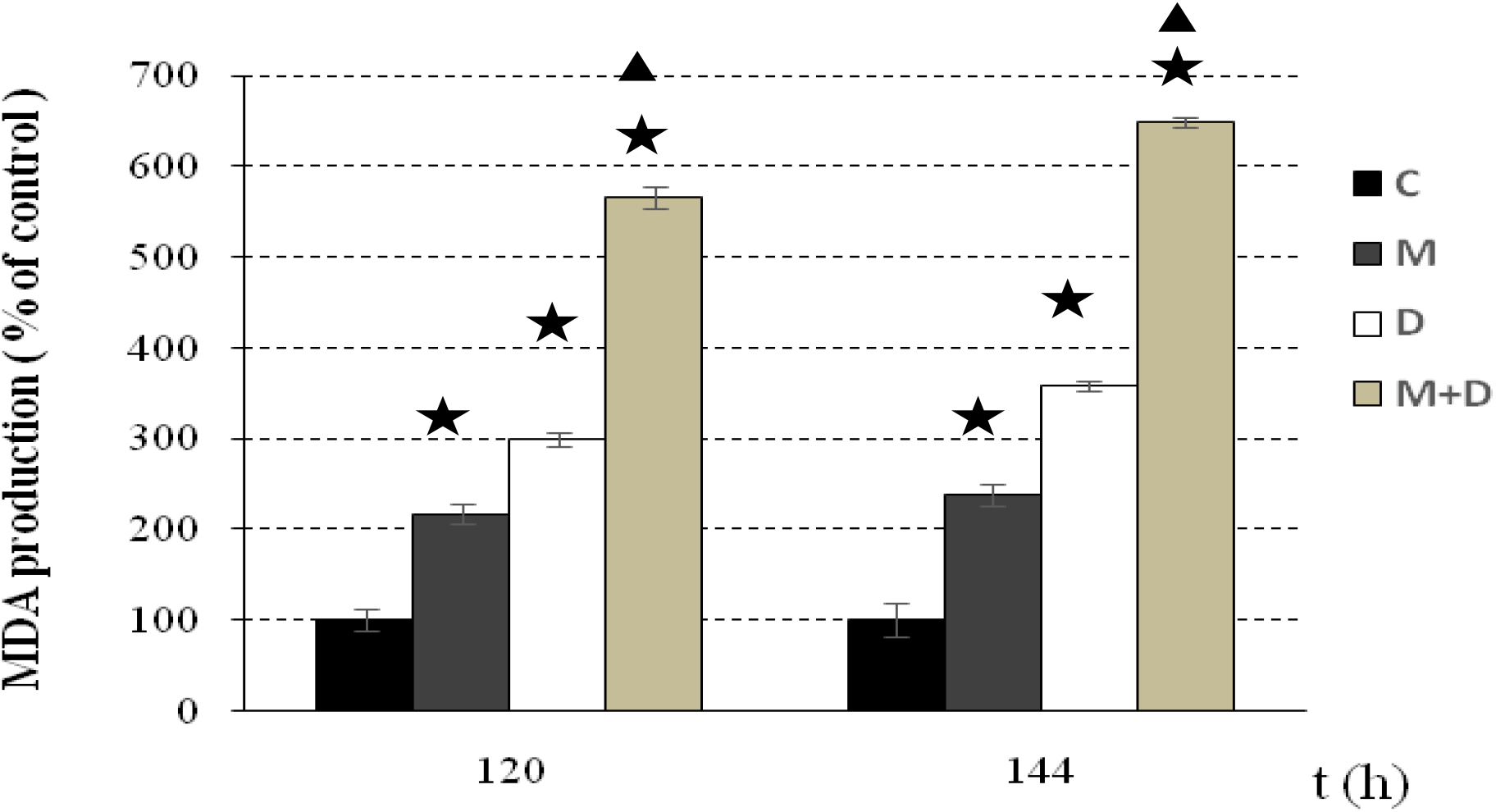
Determination of NMDA production. Values are reported as percentage of the control untreated cells, by considering it as 100%. Each value represents the mean ± SEM of four independent experiments, each done in duplicate. Star indicates the value significantly different from the control untreated cells (p < 0.05). Triangle indicates value significantly different from all the other treatments at the same time point (p < 0.05).

### Gene and protein expression (CD133, Nestin, GFAP, Notch4)

Gene expression analyzed by Real PCR technique. After EMF, TMZ and co-administration of(EMF+TMF). CD133, Nestin and Notch4 decreased but, GFAP increased. CD133 mRNA expression increased 50%,60% and 90% by exposure to EMF and TMZ alone and concomitantly after 144h. we observed the same pattern for Nestin, decreasing about 50%, 43% and 83%, 144h after applying EMF, TMZ alone and together. Also decreasing in Nestin mRNA was detected about 62%,60% and 93%by administration EMF, TMZ and both of them after 144h. conversely, over-expression of GFAP was detected about 14, 20.32 folds by exposure EMF, TMZ individually and in combination together after 144h in compare to control. We got the same result for GFAP protein expression, so expression increased 70%, 100% and 150% after treatment by EMF, TMZ alone and simultaneously up to 144h. all of these genes are involved in differentiation. Reduction of the three genes (CD133, Nestin, Notch4) and increase of GFAP indicate the beginning of the differentiation (fig. 5,6,7).

**Fig. 5.**
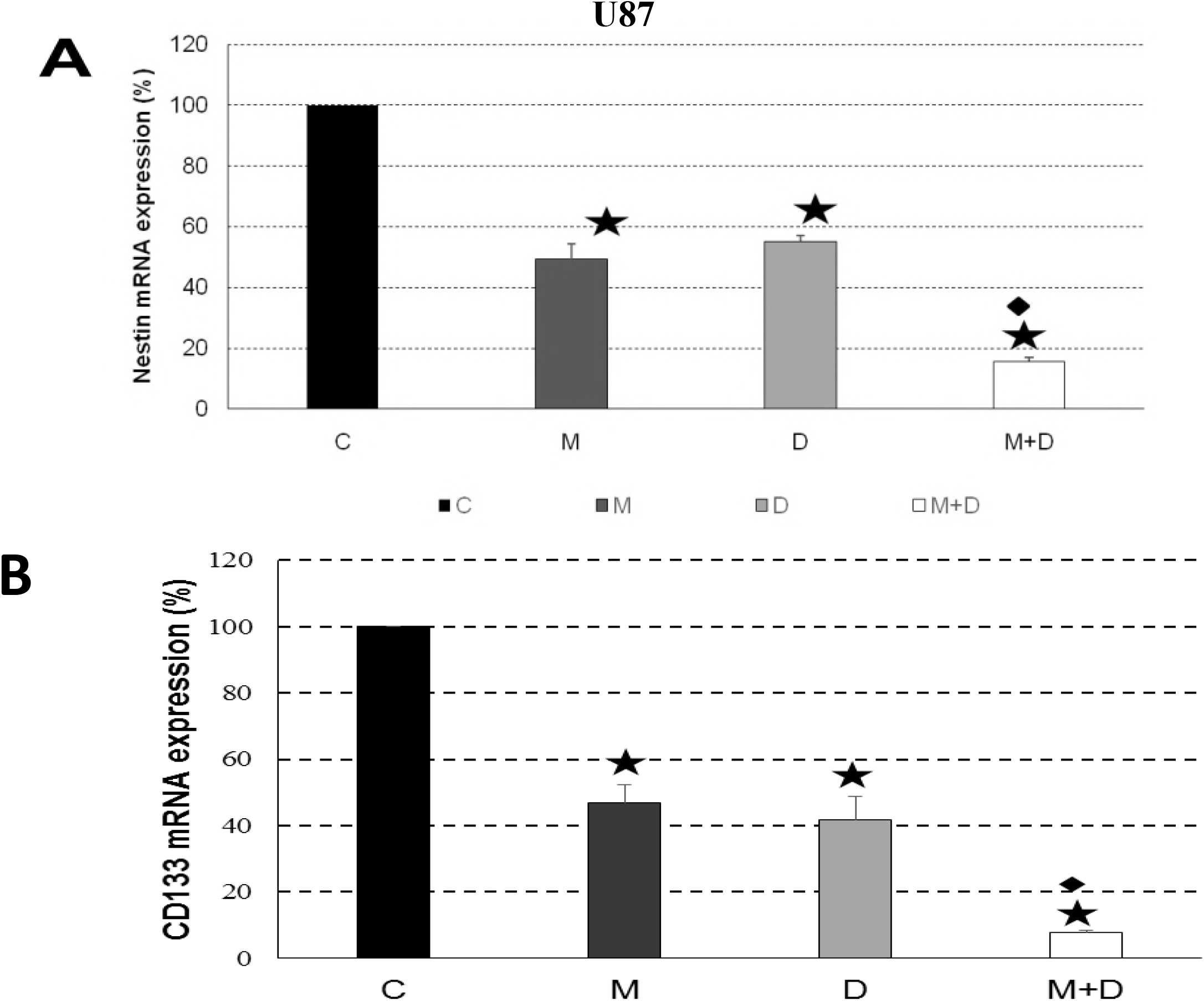
Nestin and CD133 mRNA expression levels (real-time PCR). Data are expressed as a relative value, in arbitrary units (a.u.), upon b-actin gene expression. Each value represents the mean ±SEM of four independent experiments, each done in duplicate. Star indicates the value significantly different from the control untreated cells (p < 0.05). Rhombus indicates value significantly different from all the other treatments at the same time point (p < 0.05).

**Fig. 6.**
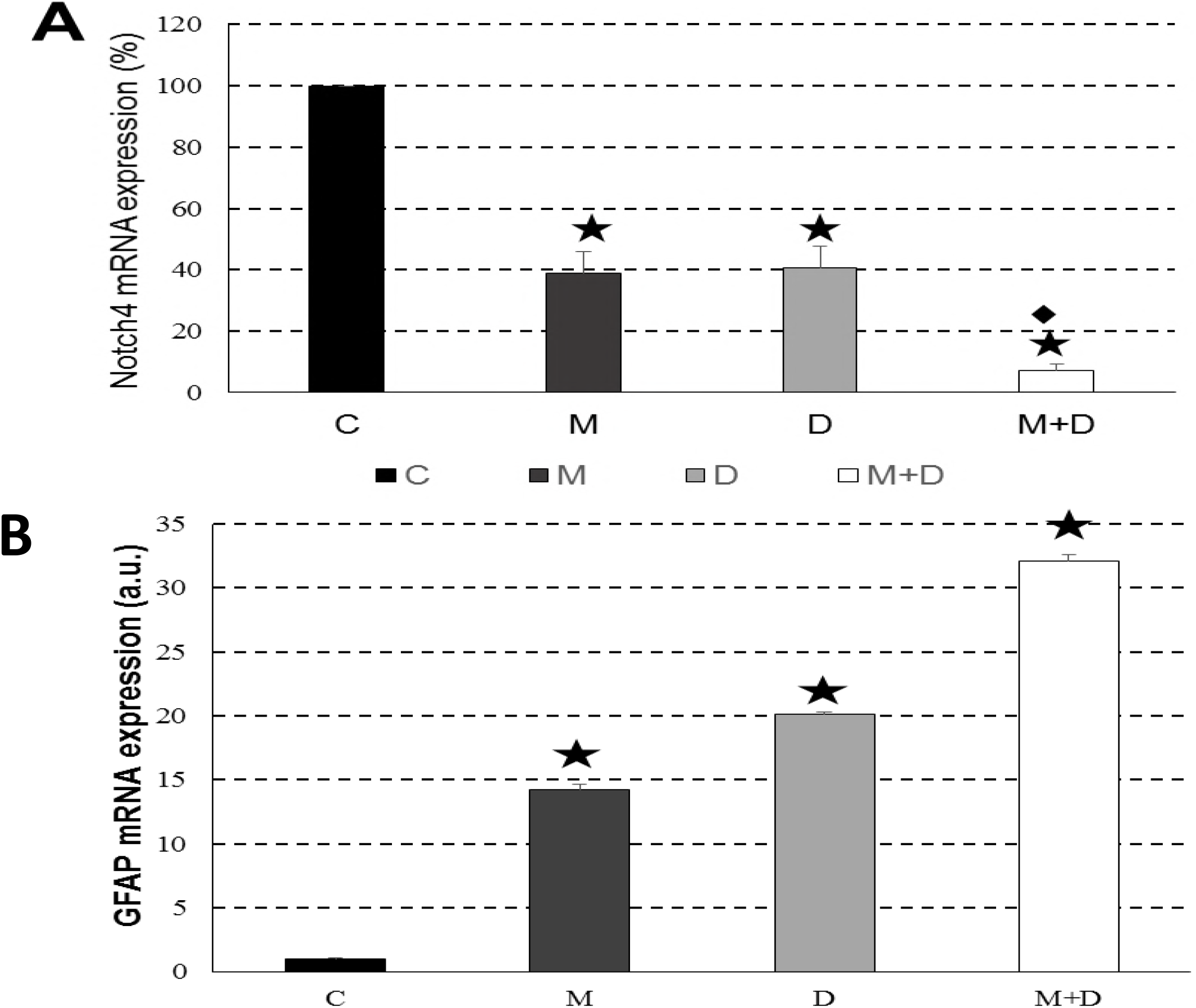
Notch4 and GFAP mRNA expression levels (real-time PCR). Data are expressed as a relative value, in arbitrary units (a.u.), upon b-actin gene expression. Each value represents the mean ±SEM of four independent experiments, each done in duplicate. Star indicates the value significantly different from the control untreated cells (p < 0.05). Rhombus indicates value significantly different from all the other treatments at the same time point (p < 0.05).

**Fig.7.**
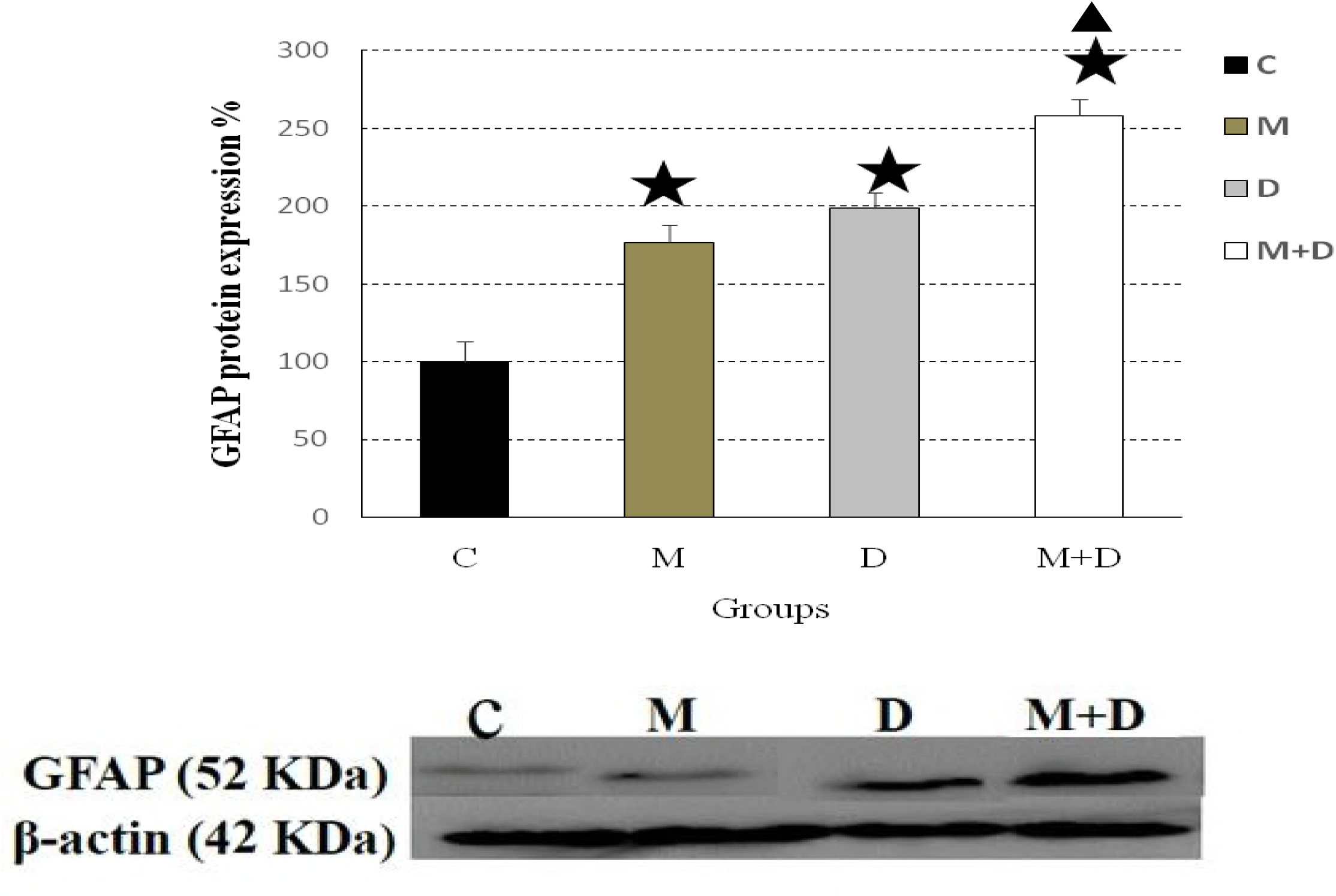
GFAP protein expression (Western blots) of U87 cells exposed to a 100 Hz, 100 G, EMF and/or treated with 100 μM TMZ for up 144 h. Values are reported with respect to the optic density (OD) of the control untreated cells, by considering it as 100%. Each value represents the mean ± SEM of four independent experiments, each done in duplicate. Star indicates the value significantly different from the control untreated cells (p < 0.05).

### Morphological changes

The morphology of U87 cells was monitored during the experiments and after144h. using light microscopy. The U87 cells grew as cultures of cells with multiple, short, fine process (neurites). Cultures grew to high density. But after exposure EMF and TMZ the number of cells decreased dramatically, in addition, EMF and TMZ induced significant morphological change in U87 cells, mainly including an elongated shape with elongated neurites, intercellular connection and neurite branching (fig.8).

**Fig.8.**
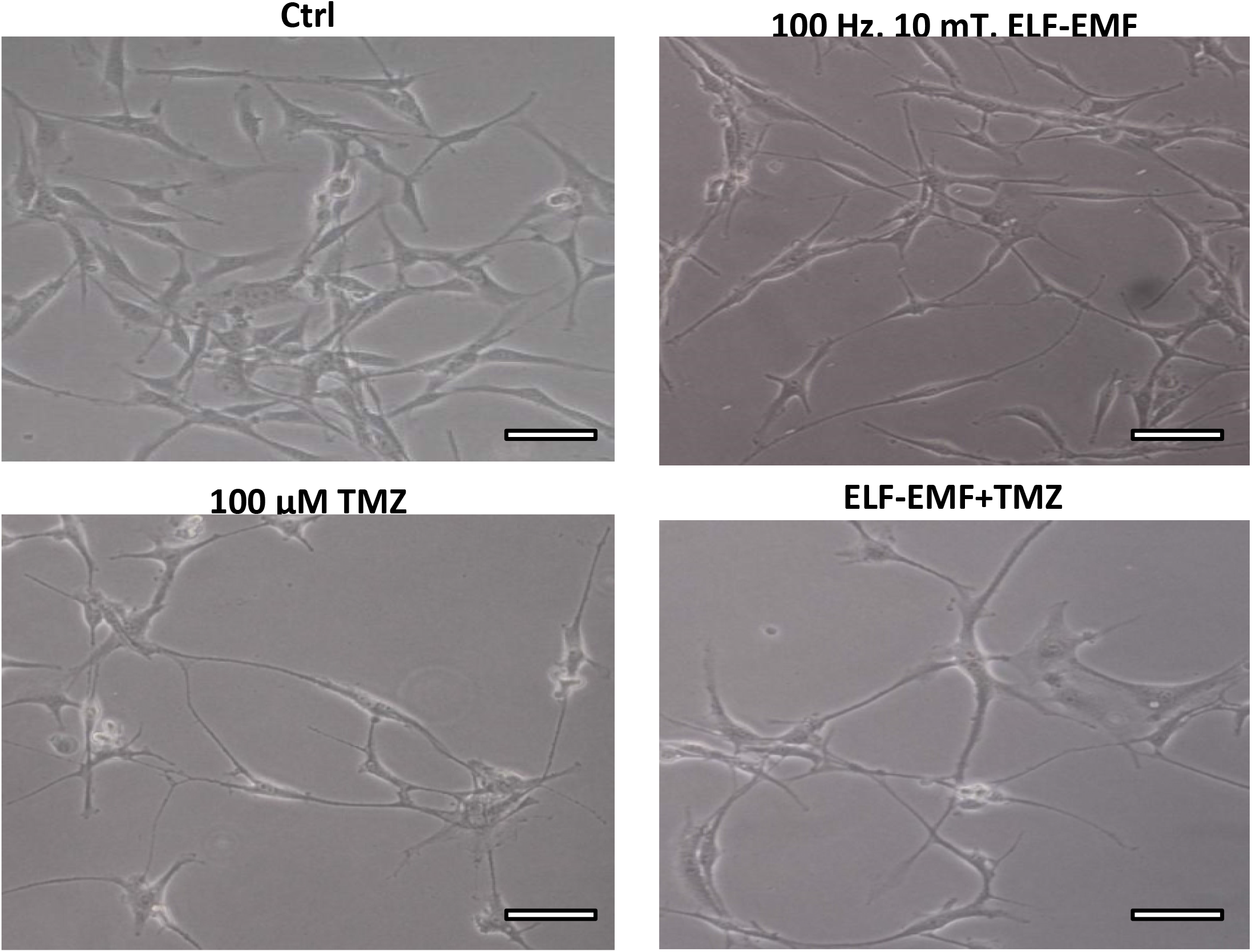
Cell morphology anlysis after exposure to a 100 Hz, 100 G, EMF and/or treated with 100 μM TMZ for up 144 h. LM micrographs of cell (U87) cultures undergone to: no treatment (Control), exposure to an EMF with a frequency of 100 Hz and an amplitude of 100 G (100 Hz,100 G, EMF), treatment with 100 μM TMZ (100 μM TMZ), or to the combined treatment (EMF + TMZ). show representative cell shape alterations. after exposure EMF and TMZ the number of cells decreased dramatically,in addition to ELF and TMZ induced significant morphological changes in U87 cells, mainly including an elongated shape with elongated neurites, intercellular connections and neurite branching. Bars = 20 μm.

## Discussion

To investigate the proliferation and differentiation phenomena induced by EMF and TMZ in u87 glioma cell line, *in vitro* model system was used to assess biological effects of exposure. As Temozolomide is a potent first choice of chemotherapy for glioblastoma patients, we evaluated the effect of this drug alone and concomitant with EMF on glioma cell line proliferation and differentiation. To find these effects, we assessed expression of Nestin and CD133 as neural stem cell marker and GFAP as astrocyte differentiation marker. Furthermore, the possible involvement of Notch signal transduction on proliferation and differentiation was also studied. Interestingly, Nestin, CD133, and Notch4 decreased, but GFAP increased. These results show that this combination therapy plays a role in differentiation of GCSCs and glioma stem cell pool reduction.

### Proliferation: (stem cell reduction) EMF and TMZ

Brain tumor initiating cells (BTICs) are implicated in the initiation, development of GBMs and radiotherapy resistance [4,32], and also have some properties of normal stem cells [33,34]. Glioma stem cells have characteristics such as self-renewal, multilinage differentiation ability and express various neural stem cell markers such as Nestin, CD133, and olig2 [35]. CD133+cells present more probability of inducing CSC than CD133^-^ do [36] and has been accepted as a standard marker of GSCs, although may not be the ideal one.

The ELF-EMF has been reported to down-regulated early neuronal marker Nestin [37].Temozolomide has also been reported to have the abilities to deplete cancer stem cell, that is dose- and time-dependent [14]. The results of our study showed that Nestin and CD133 reduced in EMF and Drug groups that were prominent in the combination group.

### Differentiation

To investigate the potential role of TMZ and EMF in differentiation, GFAP was evaluated. Since Notch is known to be implicated in glial cell fate decision, proliferation, and differentiation and concluded to be a marker of differentiated and less differentiated glioma cell [38], we also tested this gene as a differentiation marker. Glial fibrillary acidic protein as a marker of differentiated astrocytes [39], is downregulated with increasing grade of astrocytoma meaning that, GFAP is important for regulating or maintaining glial cell growth. Expression of GFAP neurofilament protein in GBM may have a lower recurrence rate and better prognosis [40]. Dell’albani et al., reported that “Notch4 increased from astrocytoma grade II to GBM” [38]. Our result showed increased GFAP expression in three group of the experiment compared to control, that was very high in magnetic and drug combined treated group. Conversely, notch4 decreased that is compatible with the result of Dell’ Albani et al.

TMZ has been reported to be involved in the differentiation process of GSCs. Villalva et al., demonstrated that GSCs engaged in the differentiation process were more sensitive to TMZ [41]. In the study of Fu J et al., freshly resected glioblastoma specimen revealed elevated expression of GFAP following TMZ treatment [42]. This was also seen in a combination of Resveratrol and TMZ [13]. A recent report has shown TMZ reduced Notch 3 expression and activation in glioma [43]. In our study in TMZ group, GFAP increased and Notch4 decreased.

EMF can affect cell proliferation and differentiation by influencing the expression of relevant genes and proteins [44]. Pulsed electromagnetic fields have been reported to promote survival and neuronal differentiation of human bone marrow mesenchymal stem cell (BM-MSCs). The electromagnetic field can induce cell differentiation of stem cells [22]. In our study EM group showed differentiation pattern.

## Mechanism

magnetic gradients can intrude into cells and act directly on cell organelles. it is still not clear how the weak MF component of EMF can induce signals in a single cell leading to migration or proliferation. This is called the problem of coupling the EMF to biological systems [44]. Many experiments on isolated cell systems point to the Ca^2^+ signaling pathway as the target of EMF in the first step of coupling [45,46].

we evaluated the role of Ca and found it, to be increased in EM and Temozolomide group which was prominent in the combination group. In this study, we considered some mechanism of differentiation that could be affected by EMF. Experimental investigations have addressed the interaction between EMF and calcium fluxes; because calcium is a principal regulator of several cellular processes and survival [47]. It results from a complex interplay between activation and inactivation of intracellular and extracellular calcium-permeable channels. These fluxes of intracellular calcium can occur as transient increases or as repetitive calcium oscillations, which both ultimately lead to altered cell activity [48]. The direct targets of ELF EMFs fields in generating non-thermal effects have not been distinctly recognized. Most of all EMF-mediated responses may directly be produced through voltage-gated calcium channel (VGCC) stimulation in the plasma membrane [49]. Despite some studies have shown that calcium signaling was not affected by low-electromagnetic fields, a systematic review with the inclusion of 42 studies, showed evidence for an association of LF EMF with internal calcium concentrations and calcium oscillation patterns [50]. Several studies suggest that ELF-EMF control ion transport through increased intracellular Ca^2+^ levels which has a modulatory role during cell differentiation. Some data shows ELF-EMF exposure increase neuronal differentiation of neural stem cells (NSCs) through increasing of Cav1-channel expression and function [51]. Calcium signaling plays an extensive role in many cellular regulatory processes including proliferation, apoptosis and gene transcription [52]. The role of calcium signaling in the cell proliferation depends on the cell and tissue types, being deeply different in tumoral cells when compared to physiological processes [53]. Calcium can also promote activation of Notch 1[54], that could be the mechanism of action on Notch4 changes in our study.

The balance between production of oxygen free radical and antioxidant in normal condition protect tissue damage against free radical. Oxidative stress is a well-known part of molecular and cellular tissue damage mechanisms [55]. Malondialdehyde (MDA) is a marker of oxidative stress and is produced during the attack of free radicals to membrane lipoproteins [56].

Superoxide dismutase plays an important role in alleviating tissue damage due to the formation of free radicals [57]. It acts as antioxidants by reducing more reactive species [58] and directly scavenging superoxide [59]. To determine whether oxidative damage occurs, and to what degree, MDA and SOD were evaluated. Our study showed that MDA was higher in the combined drug and EMF groups compared to control group, indicating the increase of oxidative stress. This increased level could be attributed to increasing ROS production and/or deficiency of antioxidant defense system. We also observed an increase in SOD activities. The results of many investigations have shown that calcium is essential for the production of ROS [60]. Increasing level of intracellular calcium is responsible for ROS-generation by the mitochondria. On the other hands, an increase in intracellular calcium concentration may be stimulated by ROS. Interactions between ROS and calcium signaling can be considered as bidirectional, wherein ROS can regulate cellular calcium signaling, while calcium signaling is essential for ROS production [61]. Simultaneously, Ca^2^+ may increase the level of superoxide dismutase (SOD) in animal cells. ROS and Ca^2^+ could be the first induced effects of ELF-EMF on biological systems [62,63]. There is a direct relationship between ROS and hem-oxygenase-1and SOD. Our previous study showed an increase in ROS, HO-1 and P53 [17].

Through redox system, over-expression of the superoxide dismutase (SODs) in vitro increases cell differentiation, decreases cell growth and proliferation and can reverse a malignant phenotype to a nonmalignant one [58,64]. It is likely that the effects of SOD on cell growth and differentiation are tightly connected to the antioxidant/oxidant balance and thus the oxidizing/reducing environment. But, in vivo, all tumors do not necessarily respond equally to the same signals. With a few exceptions, when antioxidant levels are augmented by either genetic or pharmacologic means, the impact on tumor cells both in vitro and in vivo has been shown to inhibit tumor growth and have suppressive effect on the malignant phenotype of glioma cell [65]. We found SOD and p53 were increased to combat ROS increasing. Our results seem to support the report of Ehnert et al., that extremely low-frequency electromagnetic fields cause antioxidant defense mechanisms [66].

## P53 and notch signaling

Many signaling pathways have been reported to be involved in the differentiation of glioma tumor [9,38,67,68]. P53 can regulate Notch signaling activity but Notch can also regulate p53, in reciprocal positive or negative feedback loops that are important for cell proliferation and cancer development. Positive or negative reciprocal regulation of the two pathways can vary with cell type and cancer stage [69,70]. Notch pathway can blockade and depletes stem-like cell in GBM [71]. Ca2+ can promote activation of Notch 1 and p53 [54].The role of Notch pathway in the tumorigenesis is highly variable. It can be tumor suppressive or pro-oncogenic and can either suppress or promote differentiation typically depend on the cellular context [69,72]. The results of our study and others mentioned here, regarding the mechanism of action of EMF on differentiation are summarized in (Fig.9.)

**Fig.9.**
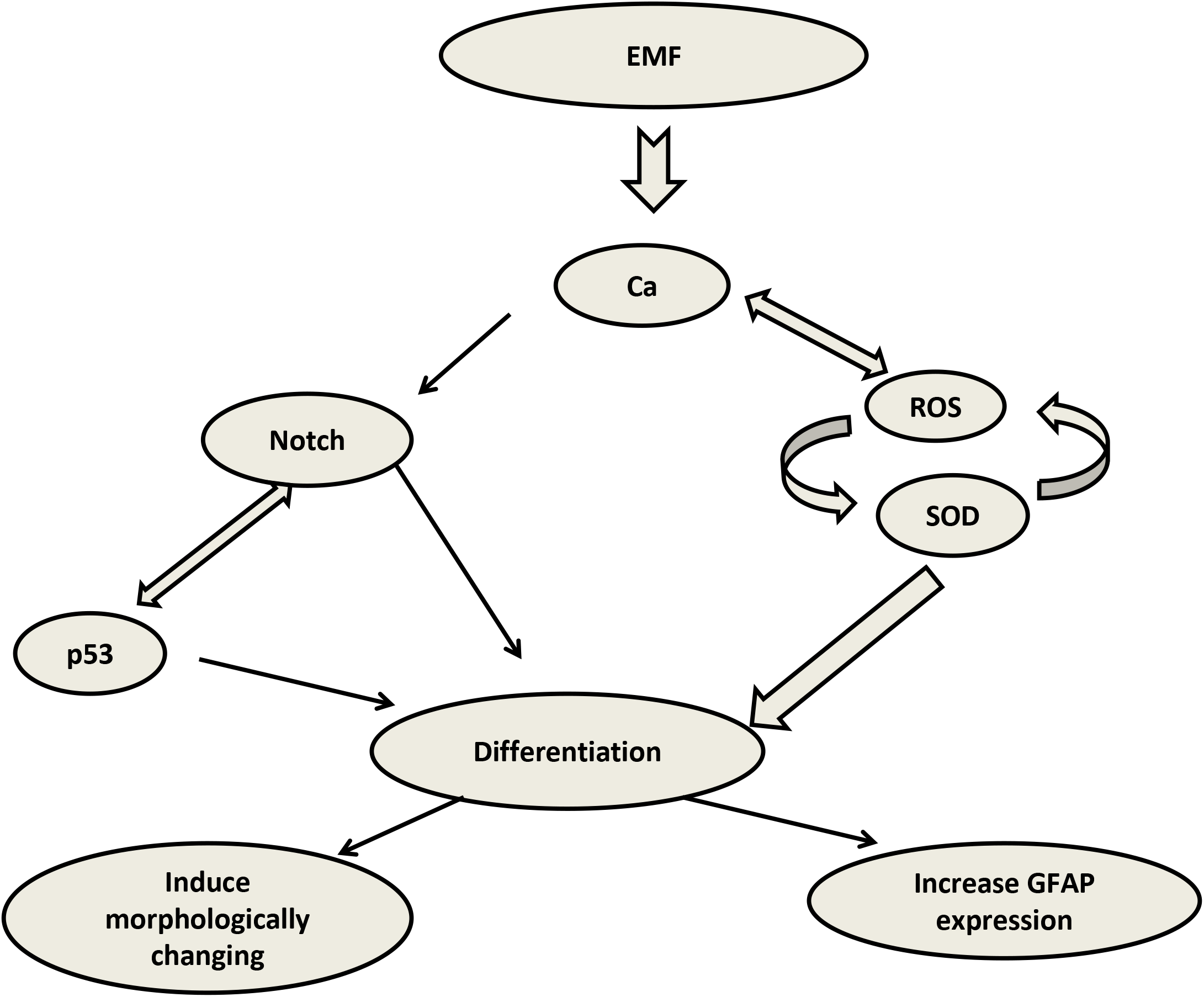
Effects of EMF on cell differentiation; calcium signaling pathway may be the first step of targeting of EMF. Calcium can promote activation of Notch4 and ROS. The interaction between ROS and SOD can be considered bidirectional. Simultaneously, calcium may increase the level of SOD. Over-expression of the SOD increase cell differentiation. P53 and SOD increase to combat ROS increasing. P53 can regulate Notch signaling but Notch can regulate P53. Both P53 and Notch increase differentiation.

## Conclusion

This study showed that EMF and TMZ treatment of glioma cell line U87 induced growth arrest as confirmed by neural progenitor/ precursor markers, Nestin, CD 133, and differentiation by GFAP. In our study, the combination of TMZ and EMFS significantly upregulate the expression of GFAP, indicating that the combined treatment could induce cell differentiation. The high Notch4 expression is associated with a less differentiated and possibly more aggressive tumor. In our study decrease of Notch4 was associated with differentiation.

The mechanism of signal transduction distal to the receptor involves intracellular pathways was due to changes in intracellular calcium. This long-term response involved persistent changes in the function of cells, decreased proliferation, changes in gene expression and differentiation through Notch, ROS, P53, and SOD. Super Oxide Dismutase acts similar to P53 as tumor suppressor genes and directly affect differentiation. Several studies have reported a relationship between tumor suppression and neural differentiation in GBM. A network of signal transduction and a complex crosstalk between all or some cell signaling are implicated in cell proliferation and differentiation of GSCs (fig. 9.).

Differentiation of tumor-initiating cells of GBM by EMF and TMZ support the new approach to GBM treatment. However, the anti-glioblastoma effects of EMF and the mechanisms underlying these activities remain to be elucidated. These findings may open new avenues for identifying a therapeutic target in the differentiation therapy of cancers. The above results suggest that our treatment induces differentiation of GBM cells and depletes the pool of tumorigenic BTICs.

## Acknowledgments

This study has been financed by the Neuroscience Research Center, Neuropharmacology Institute, Kerman University of Medical Sciences, Kerman, Iran. We also thank Mrs. Jamileh Mahdavi for her editorial assistance.

## References

1. David N. Louis, Eric C. Holland, J. Gregory Cairncross. Glioma classification: a molecular reappraisal. Am J Pathol. 2001;159: 779–786.

2. Nikolaos A Trikalinos, Young Kwok, Olga Goloubeva, Minesh Mehta, Edward Sausville. Socioeconomic status and survival in glioblastoma. Int J Clin Exp Med. 2016;9: 4131–4136.

3. Beck B, Blanpain C. Unravelling cancer stem cell potential. Nat Rev Cancer. 2013;13: 727–738. doi:10.1038/nrc3597

4. Bao S, Wu Q, McLendon RE, Hao Y, Shi Q, Hjelmeland AB, et al. Glioma stem cells promote radioresistance by preferential activation of the DNA damage response. Nature. 2006;444: 756–760. doi:10.1038/nature05236

5. Campos B, Wan F, Farhadi M, Ernst A, Zeppernick F, Tagscherer KE, et al. Differentiation Therapy Exerts Antitumor Effects on Stem-like Glioma Cells. Clin Cancer Res. 2010;16: 2715–2728. doi:10.1158/1078-0432.CCR-09-1800

6. Pierce GB. The cancer cell and its control by the embryo. Rous-Whipple Award lecture. Am J Pathol. 1983;113: 117–124.

7. Sanai N, Alvarez-Buylla A, Berger MS. Neural Stem Cells and the Origin of Gliomas. N Engl J Med. 2005;353: 811–822. doi:10.1056/NEJMra043666

8. Singh SK, Hawkins C, Clarke ID, Squire JA, Bayani J, Hide T, et al. Identification of human brain tumour initiating cells. Nature. 2004;432: 396–401. doi:10.1038/nature03128

9. Sugimoto N, Miwa S, Nakamura H, Tsuchiya H, Yachie A. Protein kinase A and Epac activation by cAMP regulates the expression of glial fibrillary acidic protein in glial cells. Arch Biol Sci. 2016;68: 795–801.

10. Terés S, Lladó V, Higuera M, Barceló-Coblijn G, Martin ML, Noguera-Salvà MA, et al. 2-Hydroxyoleate, a nontoxic membrane binding anticancer drug, induces glioma cell differentiation and autophagy. Proc Natl Acad Sci. 2012;109: 8489–8494. doi:10.1073/pnas.1118349109

11. Johnson DR, O’Neill BP. Glioblastoma survival in the United States before and during the temozolomide era. J Neurooncol. 2012;107: 359–364. doi:10.1007/s11060-011-0749-4

12. Stupp R, Mason WP, van den Bent MJ, Weller M, Fisher B, Taphoorn MJB, et al. Radiotherapy plus Concomitant and Adjuvant Temozolomide for Glioblastoma. N Engl J Med. 2005;352: 987–996. doi:10.1056/NEJMoa043330

13. Yuan Yuan, Xue Xue, Guo Ruo-Bing, Sun Xiu-Lan, Hu Gang. Resveratrol Enhances the Antitumor Effects of Temozolomide in Glioblastoma via ROS-dependent AMPK-TSC-mTOR Signaling Pathway. CNS Neurosci Ther. 2012;18: 536–546. doi:10.1111/j.1755-5949.2012.00319.x

14. Beier D, Röhrl S, Pillai DR, Schwarz S, Kunz-Schughart LA, Leukel P, et al. Temozolomide Preferentially Depletes Cancer Stem Cells in Glioblastoma. Cancer Res. 2008;68: 5706–5715. doi:10.1158/0008-5472.CAN-07-6878

15. Barbault A, Costa FP, Bottger B, Munden RF, Bomholt F, Kuster N, et al. Amplitude-modulated electromagnetic fields for the treatment of cancer: Discovery of tumor-specific frequencies and assessment of a novel therapeutic approach. J Exp Clin Cancer Res. 2009;28: 51. doi:10.1186/1756-9966-28-51

16. Vadalà Maria, Morales-Medina Julio Cesar, Vallelunga Annamaria, Palmieri Beniamino, Laurino Carmen, Iannitti Tommaso. Mechanisms and therapeutic effectiveness of pulsed electromagnetic field therapy in oncology. Cancer Med. 2016;5: 3128–3139. doi:10.1002/cam4.861

17. Akbarnejad Z, Eskandary H, Dini L, Vergallo C, Nematollahi-Mahani SN, Farsinejad A, et al. Cytotoxicity of temozolomide on human glioblastoma cells is enhanced by the concomitant exposure to an extremely low-frequency electromagnetic field (100Hz, 100G). Biomed Pharmacother. 2017;92: 254–264. doi:10.1016/j.biopha.2017.05.050

18. Luo Y, Ji X, Liu J, Li Z, Wang W, Chen W, et al. Moderate intensity static magnetic fields affect mitotic spindles and increase the antitumor efficacy of 5-FU and Taxol. Bioelectrochemistry. 2016;109: 31–40. doi:10.1016/j.bioelechem.2016.01.001

19. Storch K, Dickreuter E, Artati A, Adamski J, Cordes N. BEMER Electromagnetic Field Therapy Reduces Cancer Cell Radioresistance by Enhanced ROS Formation and Induced DNA Damage. PLOS ONE. 2016;11: e0167931. doi:10.1371/journal.pone.0167931

20. AyŞef I-G, Zafer A, Şule O, IŞil Iş-T, Kalkan T. Differentiation of K562 Cells Under ELF-EMF Applied at Different Time Courses. Electromagn Biol Med. 2010;29: 122–130. doi:10.3109/15368378.2010.502451

21. Akbarnejad Z, Eskandary H, Vergallo C, Nematollahi-Mahani SN, Dini L, Darvishzadeh-Mahani F, et al. Effects of extremely low-frequency pulsed electromagnetic fields (ELFPEMFs) on glioblastoma cells (U87). Electromagn Biol Med. 2017;36: 238–247. doi:10.1080/15368378.2016.1251452

22. Yoshie S, Ogasawara Y, Ikehata M, Ishii K, Suzuki Y, Wada K, et al. Evaluation of biological effects of intermediate frequency magnetic field on differentiation of embryonic stem cell. Toxicol Rep. 2016;3: 135–140. doi:10.1016/j.toxrep.2015.12.012

23. Wen J, Jiang S, Chen B. The effect of 100 Hz magnetic field combined with X-ray on hepatoma-implanted mice. Bioelectromagnetics. 2011;32: 322–324. doi:10.1002/bem.20646

24. Beier D, Hau P, Proescholdt M, Lohmeier A, Wischhusen J, Oefner PJ, et al. CD133+ and CD133− Glioblastoma-Derived Cancer Stem Cells Show Differential Growth Characteristics and Molecular Profiles. Cancer Res. 2007;67: 4010–4015. doi:10.1158/0008-5472.CAN-06-4180

25. Pallini R, Ricci-Vitiani L, Banna GL, Signore M, Lombardi D, Todaro M, et al. Cancer Stem Cell Analysis and Clinical Outcome in Patients with Glioblastoma Multiforme. Clin Cancer Res. 2008;14: 8205–8212. doi:10.1158/1078-0432.CCR-08-0644

26. Liu G, Yuan X, Zeng Z, Tunici P, Ng H, Abdulkadir IR, et al. Analysis of gene expression and chemoresistance of CD133+ cancer stem cells in glioblastoma. Mol Cancer. 2006;5: 67. doi:10.1186/1476-4598-5-67

27. Diehn M, Clarke MF. Cancer Stem Cells and Radiotherapy: New Insights Into Tumor Radioresistance. JNCI J Natl Cancer Inst. 2006;98: 1755–1757. doi:10.1093/jnci/djj505

28. Chomczynski P, Sacchi N. The single-step method of RNA isolation by acid guanidinium thiocyanate–phenol–chloroform extraction: twenty-something years on. Nat Protoc. 2006;1: 581–585. doi:10.1038/nprot.2006.83

29. Schmittgen TD, Livak KJ. Analyzing real-time PCR data by the comparative *C*_T_ method. Nat Protoc. 2008;3: 1101–1108. doi:10.1038/nprot.2008.73

30. Laemmli UK. Cleavage of Structural Proteins during the Assembly of the Head of Bacteriophage T4. Nature. 1970;227: 680–685. doi:10.1038/227680a0

31. Towbin H, Staehelin T, Gordon J. Electrophoretic transfer of proteins from polyacrylamide gels to nitrocellulose sheets: procedure and some applications. Proc Natl Acad Sci. 1979;76: 4350–4354. doi:10.1073/pnas.76.9.4350

32. Lee Y, Kim KH, Kim DG, Cho HJ, Kim Y, Rheey J, et al. FoxM1 Promotes Stemness and Radio-Resistance of Glioblastoma by Regulating the Master Stem Cell Regulator Sox2. PLOS ONE. 2015;10: e0137703. doi:10.1371/journal.pone.0137703

33. Dean M, Fojo T, Bates S. Tumour stem cells and drug resistance. Nat Rev Cancer. 2005;5: 275–284. doi:10.1038/nrc1590

34. Reya T, Morrison SJ, Clarke MF, Weissman IL. Stem cells, cancer, and cancer stem cells. In: Nature [Internet]. 1 Nov 2001 [cited 7 May 2018]. Available: https://www.nature.com/articles/35102167

35. Galli R, Binda E, Orfanelli U, Cipelletti B, Gritti A, Vitis SD, et al. Isolation and Characterization of Tumorigenic, Stem-like Neural Precursors from Human Glioblastoma. Cancer Res. 2004;64: 7011–7021. doi:10.1158/0008-5472.CAN-04-1364

36. Vacas-Oleas A, de la Rosa J, García-López R, Vera-Cano B, Gallo-Oller G, Alfaro-Larraya M. In vitro tumorigenicity and stemness characterization of the U87MG glioblastoma cell line based on the CD133 cancer stem cell marker. Open Access Sci Rep. 2013;2: 609.

37. Kim H-J, Jung J, Park J-H, Kim J-H, Ko K-N, Kim C-W. Extremely low-frequency electromagnetic fields induce neural differentiation in bone marrow derived mesenchymal stem cells. Exp Biol Med. 2013;238: 923–931. doi:10.1177/1535370213497173

38. Dell’Albani P, Rodolico M, Pellitteri R, Tricarichi E, Torrisi SA, D’Antoni S, et al. Differential patterns of NOTCH1–4 receptor expression are markers of glioma cell differentiation. Neuro-Oncol. 2014;16: 204–216. doi:10.1093/neuonc/not168

39. Vincenzo Bramanti, Daniele Tomassoni, Marcello Avitabile, Francesco Amenta, Roberto Avola. Biomarkers of glial cell proliferation and differentiation in culture. Front Biosci Sch Ed. 2010;2: 558–570.

40. Varlet P, Soni D, Miquel C, Roux F-X, Meder J-F, Chneiweiss H, et al. New Variants of Malignant Glioneuronal Tumors: A Clinicopathological Study of 40 Cases. Neurosurgery. 2004;55: 1377–1392. doi:10.1227/01.NEU.0000143033.36582.40

41. Villalva C, Cortes U, Wager M, Tourani J-M, Rivet P, Marquant C, et al. O6-Methylguanine-Methyltransferase (MGMT) Promoter Methylation Status in Glioma Stem-Like Cells is Correlated to Temozolomide Sensitivity Under Differentiation-Promoting Conditions. Int J Mol Sci. 2012;13: 6983–6994. doi:10.3390/ijms13066983

42. FU Jun, LIU Zhi-gang, LIU Xiao-mei, CHEN Fu, Chris Xu-rong, SHI Hong-liu, et al. Glioblastoma stem cells resistant to temozolomide-induced autophagy. Chin Med J (Engl). 2009;122: 1255–1259.

43. Chen P-H, Shen W-L, Shih C-M, Ho K-H, Cheng C-H, Lin C-W, et al. The CHAC1-inhibited Notch3 pathway is involved in temozolomide-induced glioma cytotoxicity. Neuropharmacology. 2017;116: 300–314. doi:10.1016/j.neuropharm.2016.12.011

44. Funk RHW, Monsees TK. Effects of Electromagnetic Fields on Cells: Physiological and Therapeutical Approaches and Molecular Mechanisms of Interaction. Cells Tissues Organs. 2006;182: 59–78. doi:10.1159/000093061

45. Carson JJ, Prato FS, Drost DJ, Diesbourg LD, Dixon SJ. Time-varying magnetic fields increase cytosolic free Ca2+ in HL-60 cells. Am J Physiol-Cell Physiol. 1990;259: C687–C692. doi:10.1152/ajpcell.1990.259.4.C687

46. Pessina GP, Aldinucci C, Palmi M, Sgaragli G, Benocci A, Meini A, et al. Pulsed electromagnetic fields affect the intracellular calcium concentrations in human astrocytoma cells. Bioelectromagnetics. 2001;22: 503–510. doi:10.1002/bem.79

47. Missiaen L, Robberecht W, Bosch LVD, Callewaert G, Parys JB, Wuytack F, et al. Abnormal intracellular Ca2+homeostasis and disease. Cell Calcium. 2000;28: 1–21. doi:10.1054/ceca.2000.0131

48. Smedler E, Uhlén P. Frequency decoding of calcium oscillations. Biochim Biophys Acta BBA – Gen Subj. 2014;1840: 964–969. doi:10.1016/j.bbagen.2013.11.015

49. Pall ML. Electromagnetic fields act via activation of voltage-gated calcium channels to produce beneficial or adverse effects. J Cell Mol Med. 2013;17: 958–965. doi:10.1111/jcmm.12088

50. Golbach LA, Portelli LA, Savelkoul HFJ, Terwel SR, Kuster N, de Vries RBM, et al. Calcium homeostasis and low-frequency magnetic and electric field exposure: A systematic review and meta-analysis of in vitro studies. Environ Int. 2016;92–93: 695–706. doi:10.1016/j.envint.2016.01.014

51. Piacentini R, Ripoli C, Mezzogori D, Azzena GB, Grassi C. Extremely low-frequency electromagnetic fields promote in vitro neurogenesis via upregulation of Cav1-channel activity. J Cell Physiol. 2008;215: 129–139. doi:10.1002/jcp.21293

52. Brini M, Carafoli* E. Calcium signalling: a historical account, recent developments and future perspectives. Cell Mol Life Sci CMLS. 2000;57: 354–370. doi:10.1007/PL00000698

53. Pinto MCX, Kihara AH, Goulart VAM, Tonelli FMP, Gomes KN, Ulrich H, et al. Calcium signaling and cell proliferation. Cell Signal. 2015;27: 2139–2149. doi:10.1016/j.cellsig.2015.08.006

54. Luo Xin, Jin Rui, Wang Fang, Jia Bo, Luan Kang, Cheng Feng-Wei, et al. Interleukin-15 inhibits the expression of differentiation markers induced by Ca2+ in keratinocytes. Exp Dermatol. 2016;25: 544–547. doi:10.1111/exd.12992

55. Gorrini C, Harris IS, Mak TW. Modulation of oxidative stress as an anticancer strategy. Nat Rev Drug Discov. 2013;12: 931–947. doi:10.1038/nrd4002

56. Gaweł S, Wardas M, Niedworok E, Wardas P. [Malondialdehyde (MDA) as a lipid peroxidation marker]. Wiadomosci Lek Wars Pol 1960. 2004;57: 453–455.

57. Young IS, Woodside JV. Antioxidants in health and disease. J Clin Pathol. 2001;54: 176–186. doi:10.1136/jcp.54.3.176

58. Kinnula VL, Crapo JD. Superoxide dismutases in malignant cells and human tumors. Free Radic Biol Med. 2004;36: 718–744. doi:10.1016/j.freeradbiomed.2003.12.010

59. Matés >J. M. Effects of antioxidant enzymes in the molecular control of reactive oxygen species toxicology. Toxicology. 2000;153: 83–104. doi:10.1016/S0300-483X(00)00306-1

60. Görlach A, Bertram K, Hudecova S, Krizanova O. Calcium and ROS: A mutual interplay. Redox Biol. 2015;6: 260–271. doi:10.1016/j.redox.2015.08.010

61. Gordeeva AV, Zvyagilskaya RA, Labas YA. Cross-Talk between Reactive Oxygen Species and Calcium in Living Cells. Biochem Mosc. 2003;68: 1077–1080. doi:10.1023/A:1026398310003

62. Falone S, Grossi MR, Cinque B, D’Angelo B, Tettamanti E, Cimini A, et al. Fifty hertz extremely low-frequency electromagnetic field causes changes in redox and differentiative status in neuroblastoma cells. Int J Biochem Cell Biol. 2007;39: 2093–2106. doi:10.1016/j.biocel.2007.06.001

63. Morabito C, Guarnieri S, Fanò G, Mariggiò MA. Effects of Acute and Chronic Low Frequency Electromagnetic Field Exposure on PC12 Cells during Neuronal Differentiation. Cell Physiol Biochem. 2010;26: 947–958. doi:10.1159/000324003

64. Zhong W, Oberley LW, Oberley TD, Clair DKS. Suppression of the malignant phenotype of human glioma cells by overexpression of manganese superoxide dismutase. Oncogene. 1997;14: 481–490. doi:10.1038/sj.onc.1200852

65. Zhang Y, Zhao W, Zhang HJ, Domann FE, Oberley LW. Overexpression of Copper Zinc Superoxide Dismutase Suppresses Human Glioma Cell Growth. Cancer Res. 2002;62: 1205–1212.

66. Ehnert S, Fentz A-K, Schreiner A, Birk J, Wilbrand B, Ziegler P, et al. Extremely low frequency pulsed electromagnetic fields cause antioxidative defense mechanisms in human osteoblasts via induction of •O 2 – and H 2 O 2. Sci Rep. 2017;7: 14544. doi:10.1038/s41598-017-14983-9

67. Sareddy GR, Viswanadhapalli S, Surapaneni P, Suzuki T, Brenner A, Vadlamudi RK. Novel KDM1A inhibitors induce differentiation and apoptosis of glioma stem cells via unfolded protein response pathway. Oncogene. 2017;36: 2423–2434. doi:10.1038/onc.2016.395

68. Sato Atsushi, Sunayama Jun, Matsuda Ken-ichiro, Seino Shizuka, Suzuki Kaori, Watanabe Eriko, et al. MEK-ERK Signaling Dictates DNA-Repairf Gene MGMT Expression and Temozolomide Resistance of Stem-Like Glioblastoma Cells via the MDM2-p53 Axis. STEM CELLS. 2011;29: 1942–1951. doi:10.1002/stem.753

69. Dotto GP. Crosstalk of Notch with p53 and p63 in cancer growth control. Nat Rev Cancer. 2009;9: 587–595. doi:10.1038/nrc2675

70. Yun Jieun, Espinoza Ingrid, Pannuti Antonio, Romero Damian, Martinez Luis, Caskey Mary, et al. p53 Modulates Notch Signaling in MCF-7 Breast Cancer Cells by Associating With the Notch Transcriptional Complex Via MAML1. J Cell Physiol. 2015;230: 3115–3127. doi:10.1002/jcp.25052

71. Fan Xing, Khaki Leila, Zhu Thant S., Soules Mary E., Talsma Caroline E., Gul Naheed, et al. NOTCH Pathway Blockade Depletes CD133-Positivef Glioblastoma Cells and Inhibits Growth of Tumor Neurospheres and Xenografts. STEM CELLS. 2009;28: 5–16. doi:10.1002/stem.254

72. Zhang Min, Biswas Sangita, Qin Xin, Gong Wenrong, Deng Wenbing, Yu Hongjun. Does Notch play a tumor suppressor role across diverse squamous cell carcinomas? Cancer Med. 2016;5: 2048–2060. doi:10.1002/cam4.731

